# CNN based Heuristic Function for A* Pathfinding Algorithm: Using Spatial Vector Data to Reconstruct Smooth and Natural Looking Plant Roots

**DOI:** 10.1101/2021.08.17.456626

**Authors:** Robail Yasrab, Michael P Pound

**Affiliations:** Institute of Biomedical Engineering, Department of Engineering Science, University of Oxford, Oxford, OX3 7DQ, UK; Lab of Computer Vision, School of Computer Science, University of Nottingham, NG8 1BB, UK

**Keywords:** Deep learning, Plant Phenotyping, Dijkstra, A* Search, Heuristic Function

## Abstract

In this work we propose an extension to recent methods for the reconstruction of root architectures in 2-dimensions. Recent methods for the automatic root analysis have proposed deep learned segmentation of root images followed by path finding such as Dijkstra’s algorithm to reconstruct root topology. These approaches assume that roots are separate, and that a shortest path within the image foreground represents a reliable reconstruction of the underlying root structure. This approach is prone to error where roots grow in close proximity, with path finding algorithms prone to taking “short cuts” and overlapping much of the root material. Here we extend these methods to also consider root angle, allowing a more informed shortest path search that disambiguates roots growing close together. We adapt a CNN architecture to also predict the angle of root material at each foreground position, and utilise this additional information within shortest path searchers to improve root reconstruction. Our results show an improved ability to separate clustered roots.

## 1 Introduction

Heuristic-based pathfinding algorithms have been designed to solve various real-world challenges. From road network planning, mobile navigation, to network packet routing, these algorithms underpin many systems. Within plant phenotyping, they have found a role in some techniques aimed at reconstruction root systems [REF ROOTNAV ROOTNAV2 and others]. Deep learned semantic segmentation is effective at labelling root pixels within images, but additional processing is often required to determine how many roots exist, where these exist, and the overall topology of the root architurcture. For example a segmented image will tell us how many pixels of second order roots exist, but not how many second order roots exist, or to which first order root they are attached.

Recent work [1] has proposed combining semantic segmentation and heatmap regression with shortest path algorithms to completely reconstruct root architectures. An encoder-decoder network is used to identify and classify foreground pixels and their root type (first or second order) as well as the probable locations of the seed and root tips. Dijkstra’s algorithm [ref] and A* heuristic search [2] are used to track the likely path of roots through the image between tips and the seed, with the search weighted based on the output of the segmentation. A higher weight penalty is applied to roots of a different class, and background, causing paths to generally follow root material correctly. The drawback of this approach, and what we propose to address with this work, is that the use of simple pixel class to weight a shortest path search inevitably leads to ambiguity where roots meet or cross. When two roots travel in close proximity causing paths to reach the same position, the heuristic searches for both roots are guaranteed to produce identical paths from that point onwards, as one of the segmented roots will present a shorter path to the goal. This problem becomes more noticeable on dense root systems where close proximity is more common. In this work we present an adapted heuristic search that considers both semantic segmentation output, and predicted root angle at each position. We extend the encoder-decoder architecture used in [1] to predict root angle at each pixel by encoding angle into separate X and Y channels of a unit vector. The CNN directly regresses these components, from which we can decode the angle during inference. We use an adapted MSE loss to train the network, which we term Masked MSE, that applies no loss for background pixels. Figure 1 provides an overview of this network architecture. We then extend traditional A* search with an additional weight function that penalises paths travelling perpendicular to the direction of the root material. We show that this adapted search provides more reliable root reconstruction where roots travel close together, by preventing paths crossing other to other root material. The approach also provides a slight performance improvement, as the search explores fewer neighbouring pixels perpendicular to the true root angle. This work makes the following contributions:

1. Direct prediction or per-pixel root angle using a CNN. Angle is encoded as a unit vector and predicted by two additional channels at the network output. To our knowledge this is the first work to directly regress angle measurements in this way.
2. Use of a masked loss function to focus learning on root pixels, ignoring background where angle is irrelevant
3. An extension to traditional Dijkstra and A* heuristic search that incorporates weight penalties for directions of travel that contradict the regressed angle measurements.
4. Our analysis shows that our approach provides better root system coverage than a traditional shortest path search, as well as a performance improvement.

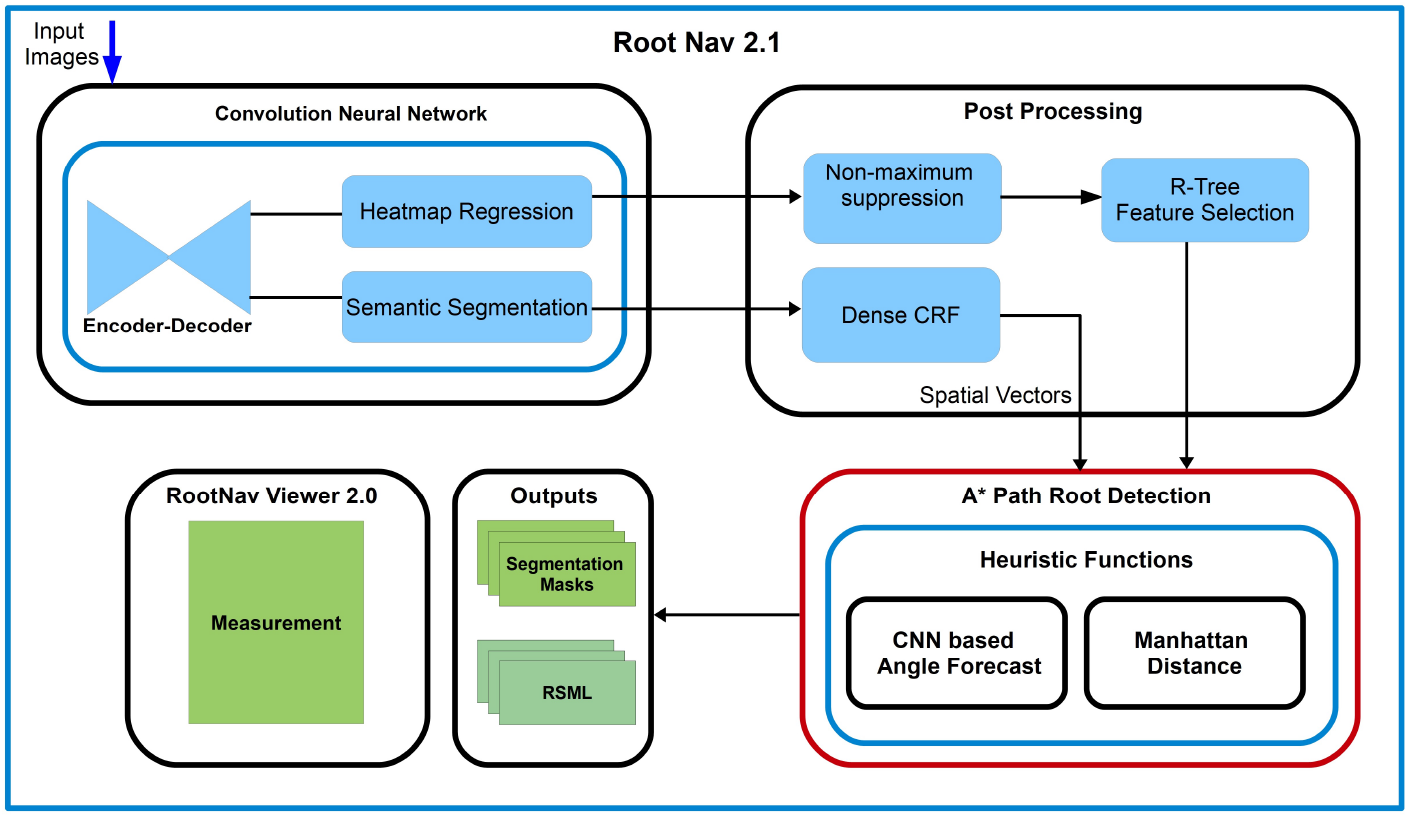

**Fig. 1.**
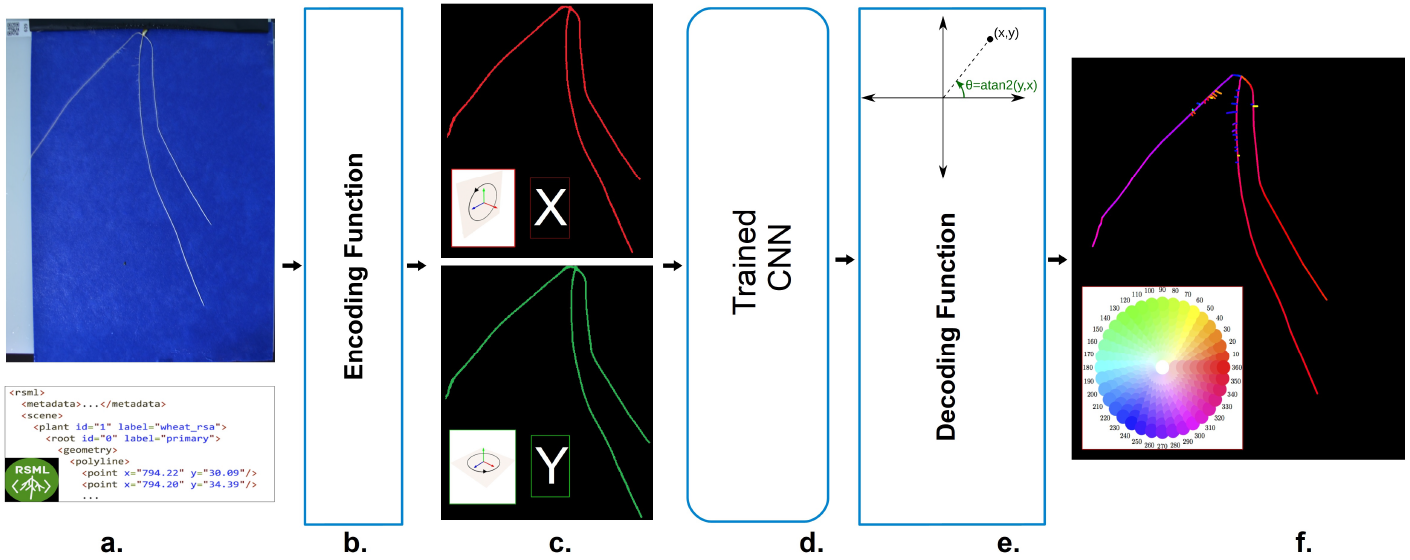
Unit Vector Encoding, Training and Decoding as Directional Angle: **a.** Input data composed of images and corresponding annotations in RSML format. **b.** Encoding function that encodes spatial vector details into trainable parameters. **c.** X and Y spatial vector represented as two trainable feature maps. **d.** A custom-designed CNN network designed to deal with different inputs and annotations and produce diverse segmentation and classification outcomes.. **e.** A decoding function that transforms a trained spatial vector to root growth angles. **f.** A visual depiction of the forecast of the directional angle using the HSV color wheel.

The paper is structured as follows. Section 2 provides an overview of the previous work in this area. Section 3 offer a description of data, deep network training and shortest path approach. The results and analysis 4 section will provide a comprehensive review of the results.

## 2 Background

Plant phenotyping plays a pivotal role in plant science research, enabling large-scale genetic discovery, and the breeding of more resilient traits [3]. Advances in plant phenotyping are therefore a key contributor to the push for global food security, particularly in the face of climate change. Root phenotyping focuses on the analysis of root growth, and has become a widely used tool to explore the influence of abiotic stresses such as high temperate and drought on a plant’s ability to take up water and nutrients [4]. Many works exist to quantify root architectures both in 2D images [5,6] and 3D. Root imaging represents a significant challenge, with highly variable structure between species and imaging modality. Recently, the move towards deep learning solutions to this task has seen a number of notable improvements in root segmentation and reconstruction techniques such as in-painting to correct problematic images. While the performance of deep networks on root segmentation problems is impressive, quantifying the resulting root architecture remains challenging. Without knowledge of which root pixels comprise a single root organ, it is difficult to measure lengths, counts, angles, or other key measures used in genetic studies. Many approaches extend segmentation with post-processing that extracts root architecture. Recent work in this area, [1], proposed to use the segmentation and heatmap regression output of a CNN to drive a series of shortest paths searches throughout the image. Similar to a previous tool [7], a shortest path algorithm such as A* search is used to find an optimal path between a root tip, and the source of that same root (either a seed, or another higher-order root). The search is performed on an 8-way connected graph, initialised using the segmented output of the CNN, with foreground root pixels assigned a smaller (preferable) weight to background pixels. This search is conducted separately for each root, before the extracted topology is encoded in the common XML-based RSML format [8].

**Fig. 2.**
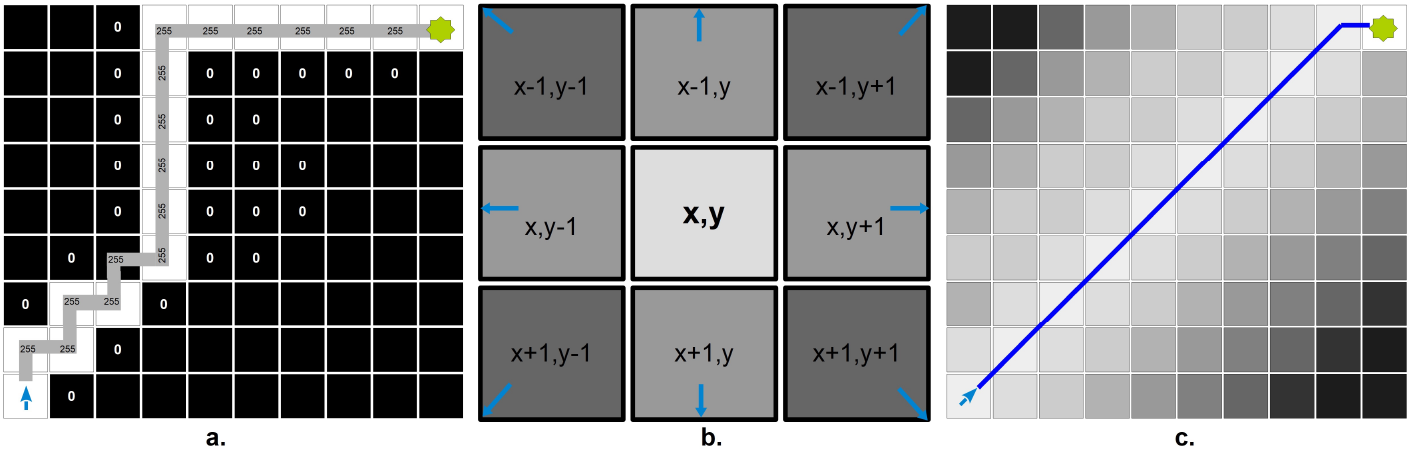
Methods Used for roots navigation and reconstruction: **a. Solving Maze:** We used the maze solving approach to navigate through the roots system. We used CNN generated GT as a maze where background pixels are no travel region where only root paths are navigation routes. **b. Manhattan distance** we have used Manhattan distance as a heuristic measure to calculate the remaining distance to the goal. We used per-pixel 8-way search to calculate the next pixel jump using *f*(*p*) = *g*(*p*) + *h*(*p*), where *g*(*p*) is the sum of all weights to *p*, and h(*p*) is the remaining distance, which we calculate as the manhattan distance, or *L*^1^-norm. **c. Weight Matrix:** The network output converted into a weight matrix, that offers a more natural pixel distribution and as a result, A* algorithm constructs a more smooth path.

The benefits of shortest path approaches like this include efficiency, and the ability to overcome gaps and mistakes in segmentation. However, care must be taken to ensure that the weight and heuristic functions used are representative of the underlying problem to be solved. In cases where root systems are dense and roots appear in close proximity, a distinction between foreground and background is not sufficient to disambiguate between the nearby paths of two different roots. This problem manifests itself as roots clustering together around single paths, rather than travelling separately where they originally appear in the image.

In this work we extend this approach to consider the angle of the roots within the image, providing additional information for a shortest path search that allows it to better separate roots. There are few comparable works on traversing images in this way in the literature, but our work is inspired by works within route planning that penalise specific turns [9] [10], and work to reconstruct arterial vessels using shortest paths with curvature regularisation [11]. Our contribution differs as our weights do not consider the current direction of travel or a measure of curvature during a search, but require only the CNN output that indicates an optimal direction of travel at each pixel. The CNN output itself is regressed using a two-channel unit vector encoding of root angle. This approach is most similar to that seen within the optical flow literature [12], in which flow fields are predicted using two-channel output. We do not use a multi-image input, and rather than optical flow choose to regress the orientation at each pixel.

## 3 Method

### 3.1 Data and Ground Truth Preparation

We make use of the dataset released in [1]. As we aim to use predicted angle to better separate root material in cases where the underlying segmentation might be ambiguous, we train and test on the Rapeseed (Brassica napus) images. These represent the most challenging within this dataset, containing the greatest number of lateral roots that also appear close together. The images are X by Y pixels in size, which we resize to 1024^2^ for training and evaluation. The dataset annotations are provided in an XML tree structure, with a hierarchy of roots represented as polylines.

As in [1], first and second order roots are rendered into separate ground truth masks using lines of thickness 4 pixels. We chose this amount as a compromise between reasonable separation between nearby roots, and the power of the network to resolve fine detail. The seed location, first and second order root tips are stored as separate lists and used to render heatmaps for regression. Heatmaps are rendered by during training where each tip or seed location is represented by using Gaussian kernels of standard deviation 2 pixels.

We extend the approach of [1] by encoding the angle at each position along each root. Since each root is represented as a polyline, the angle is identical for any position along any given line segment rather than a continuous function over the root length. We thus render line segments individually with angles calculated for each. A given line segment is represented as two points (*x*_0_, *y*_0_) and (*x*_1_, *y*_1_), from which we calculate the vector **υ** = 〈*x*_1_ — *x*_0_, *y*_1_ — *y*_0_〉, before normalising to unit vector 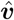. We render the x and y components of this unit vector separately into two additional ground truth masks. Each line segment is rendered with the same thickness and position as in the segmentation masks, as we will later apply a masked loss function to these pixels specifically. No angle is calculated or used for background pixels.

During inference it is straightforward to decode the x and y components for a pixel i back into a rotational angle *θ*:

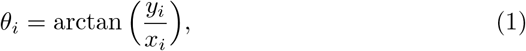

where –π ≤ *θ* ≤ π, and an angle of 0 represents the vector (0,1). Figure 3 shows a visualisation of ground truth angle for a wheat image using the HSV colour space. We next convert *θ* to an appropriate orientation for use in guiding shortest path searches. Since we prioritise paths that travel parallel to root direction rather than orthogonal to it, we elected to remap angles into the range 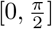 representing a range from horizontal to vertical.

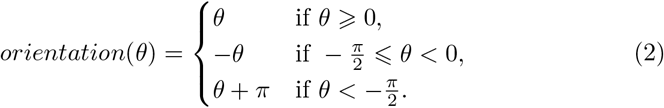

**Fig. 3.**
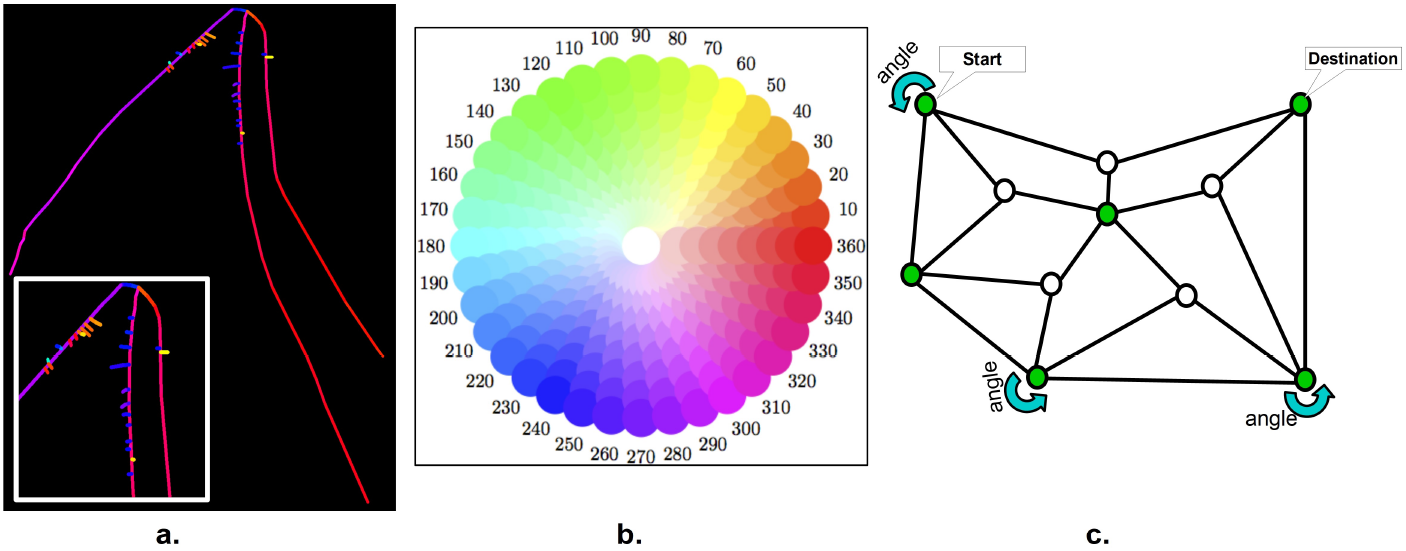
**a.** A color-based visualization of CNN based heuristic function’s output in the form of roots angle of growth, where each color represents a different angle or direction of root growth. **b.** HSV color wheel is shown for reference of colors in Fig 6 a. **c.** A visual description, how an A* path search-finding algorithm can make use of angle based heuristics to navigate a complex roots network. The angle heuristic will help the algorithm to choose the next pixel that is most suitable at each step.

Below we describe an adapted shortest path approach where this orientation can be compared against the relative movement orientation towards neighbouring pixels.

### 3.2 CNN Design and Network architecture

The proposed CNN architecture is adapted from [1], which uses an encoder-decoder architecture with separate branches for output of segmentation masks and heatmap regression of key points. We add an additional output path to return two channels representing the x and y components of the angle of the root at each pixel. An illustration of the architecture may be seen in [FIGURE REF]. We found that using separate branches for each output rather than a single layer improved performance on each sub-task, but that only a few 1×1 convolutional layers were necessary to obtain the desired accuracy. The network operates on an input of 1024^2^ pixels, with initial fast down-sampling to bring the feature maps into a practical size. For efficiency, we perform one less deconvolution operation in the decoder, meaning the output resolution of the network is 512^2^.

### 3.3 Loss Function

As described above the output of the proposed network is divided into three paths with different objectives. The first and second outputs are responsible for learning to predict the plant root segmentation and key point regression tasks.

Cross-entropy loss and mean squared error loss used to extract 2D binary outputs for segmentation tasks. We have used the same dual loss approach as of our earlier research work [1]. In this proposed system, training the CNN with traditional single loss function is not possible. We have included a special loss calculation method, “Means square Error (MMSE)” loss. So, the proposed CNN training makes us of Cross-entropy loss, mean squared error loss and Masked Means square Error (MMSE) loss for the segmentation of root and forecast of growth angle. MASKED MSE [14] loss function application for training normalized spatial vectors leads to successful results.

The training growth angle is a delicate and precise process and needs a special loss function to deal with this problem. We cant use traditional loss function as most of the values in the training mask for angle training are below zero. Therefore, using a traditional loss function will not able to train and classify significant areas of interest. Masked Means square Error or MMSE loss offers the capability to learn all areas of interest by ignoring background noise. In this special training process for angle GT, the background values set to zero, and foreground values (normalized vectors) set between 1-0. MMSE ignore all background 0 values and learn on the foreground angle of roots growth. Therefore, when we calculate and optimize the growth angle loss during backward propagation, all the background zero values get ignored at the input layer. This loss function helped us to learn exact angle values those later extracted and used as a heuristic function in the root growth redrawing process. The Eq. 3 shows the mathematical expression of the MMSE loss function:

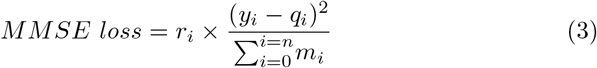

Where, y_i_ is the original ground truth, and q_i_ is the CNN output. To ignore the 0 value during loss calculation process, r_i_ mask value is used such that r_i_ = 0 if y_i_ = 0 and r_i_ = 1 otherwise. In special cases, mean normalization parameter could be enabled, so the rating value 0 could not be ignored, in this situation, r_i_ = 1 for all y_i_ is a fixed setting.

**Fig. 4.**
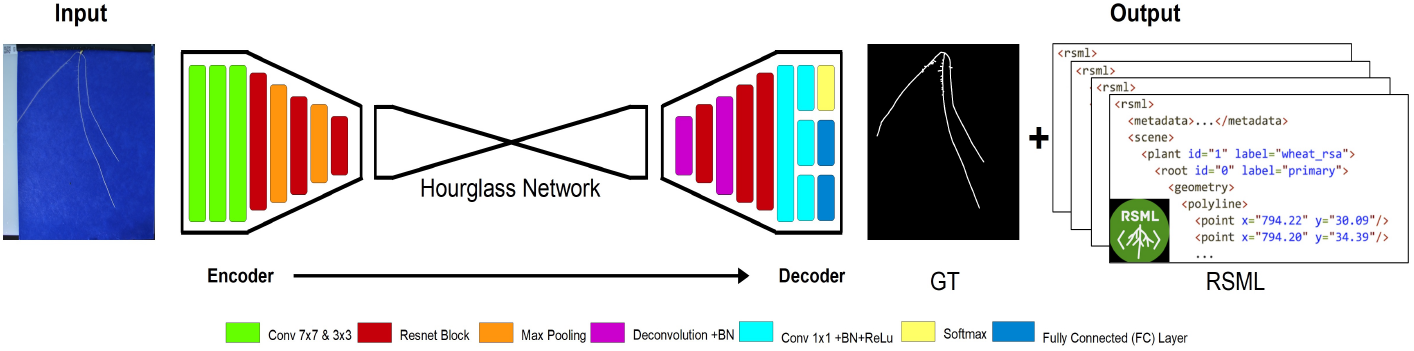
The proposed CNN Architecture: The CNN inspired by RootNav 2.0 [1] reserch. With some additional tweaks, it is capable of processing spatial vector data and also output additional feature maps to forecast angle of growth. It is a nested encoder-decoder (ED) architecture. That sandwich an hourglass CNN network [13] with a SegNet style ED. The input is processed through the encoder and upsampled in the decoder section using a series of learned deconvolutional layers upsample back to 50% of the original input size. In the end, the network split into three outputs: (i) semantic segmentation of root system (Primary root, Lateral root, Seed Location), (ii) heat map regression of root tips and finally, (iii) root growth patterns in the form of angles.

### 3.4 Training

The proposed CNN trained end-to-end using the transfer learning method on the given dataset. We have used a pre-trained CNN, that was previously trained out earlier project [1] for 500,000 iterations. The trained model is used to transfer-learn the new features in much efficient ways than training from scratch. Training the proposed CNN from scratch was not offering better results. The transfer-learning based training performed for almost 50,000 iterations using the “rm-sprop” optimizer.

The initial learning rate was set to 1*e*^-4^ and reduced by a factor of 10 after 50,000 iterations with a batch size of 6. Some data augmentation methods also used to reduce the possibility of overfitting and take advantage of better learning. Random horizontal flipping used as an augmentation method with a 50% probability during training.

## 4 Results and Analysis

This section will present a quantitative analysis of the proposed method. To ensure the method’s effectiveness, we have performed a visual and time-based analysis of the proposed method.

**Fig. 5.**
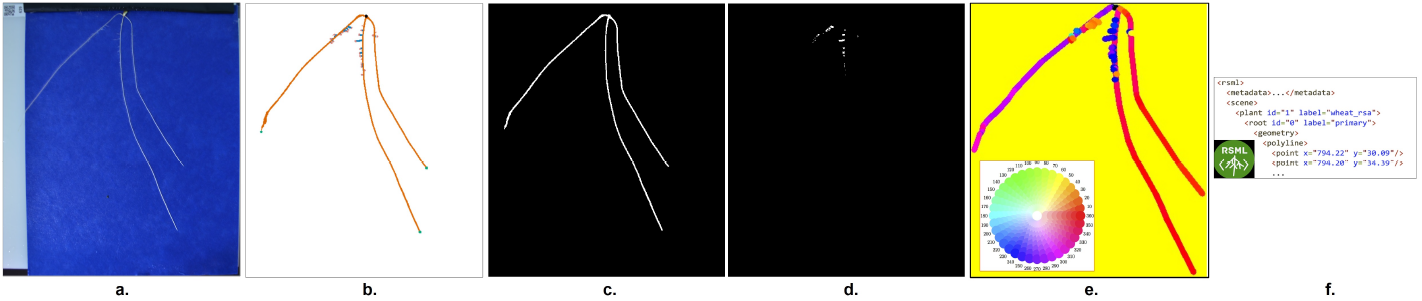
Graphical Analysis of Results: **(a)** Input Image. **(b)** Colour-coded segmentation mask. **(c, d)** Binary segmentation masks for first- and second-order roots. **(e)** Color-coded angle forecast mask represented through the HSV Colour wheel. **(f)** An RSML output that contains the entire architecture of the root system.

The Fig 6 show a side by side comparison of the effectiveness of the method. We can see that root detected by RootNav2 drawn in Fig. 6(a) looks more occluded, and most of the roots took a shorter path besides choosing the real path of root reconstruction. In this scenario, the resultant roots reconstruction looks unnatural and occluded. In the Fig 6(b), we can see that roots take the real path back to the primary source and offered a better visual reconstruction of origins. The predicted root angle assisted the AStar algorithm of the right root track, and resultant roots look more natural.

**Fig. 6.**
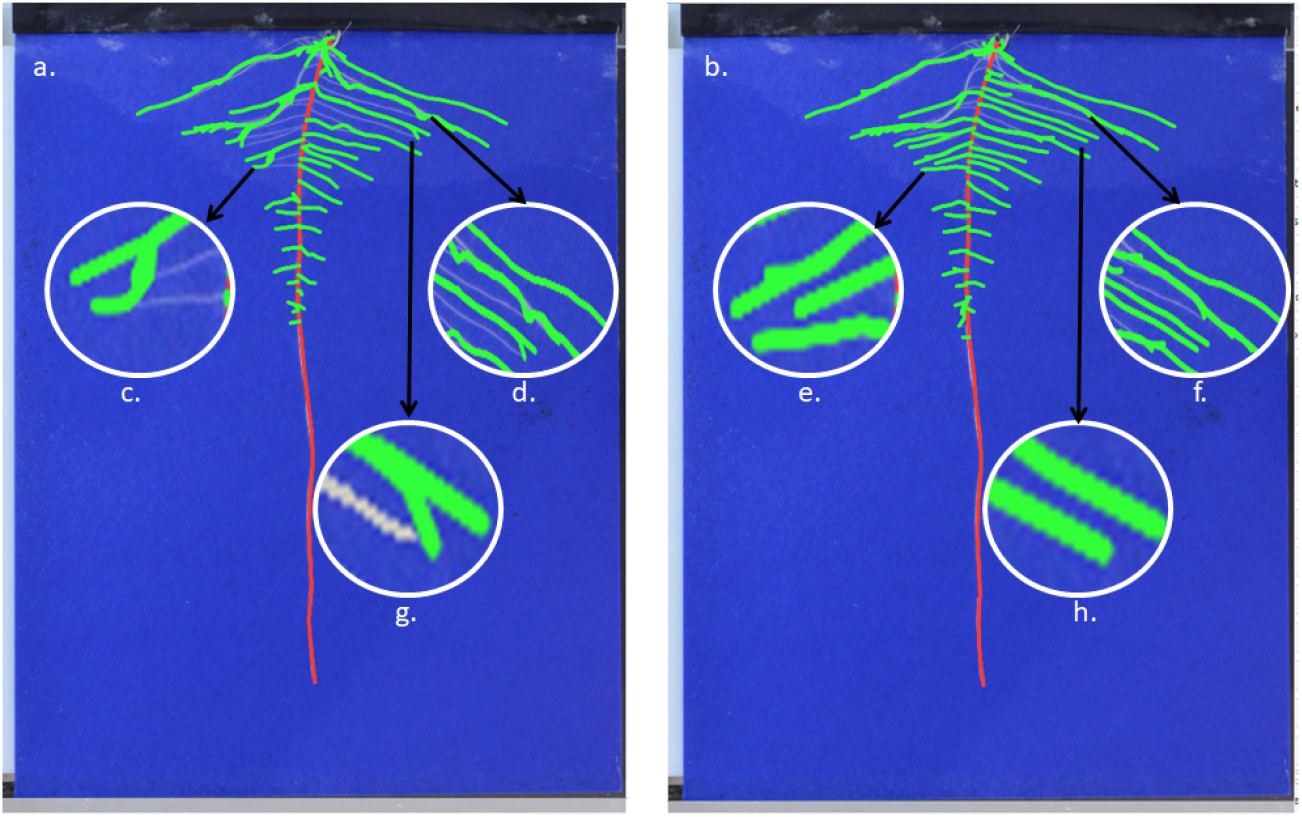
RootNav 2 Original vs. RootNav 2 with Angle forecast: **a)** Image processed through Original version of RootNav 2, b) Image processed through angle forecast version of RootNav2, **c&g)** The Astar algorithm search for the shortest path to reach back to the source of the root. In this situation, i skip the original root lane and travel through a short path; that results in an occluded and unnational roots outlook. **e&h)** the same roots mentioned earlier taking the right track during root reconstruction with additional help for angle forecast. **d)** The root reconstructed results in a jagged line that is an unnatural look for roots. **f)** with angle forecast support, RootNav2 produced more natural-looking roots.

The second analysis performed a time-based analysis to ensure the high performance of the proposed method (Table. 1). The proposed new root angle forecast method helped the ASatr algorithm better navigate the root skeleton for the root reconstruction task. We have used an 8-way manhattan distance method to search and reconstruct the root path. However, due to the angle forecast, this 8-way pathfinding gets better. The newly proposed method cuts down the additional search directions and propels the AStar on the right track. As a result, he can cut down extra search time that results in lower inference time per image. The Graph shows a time-based analysis of test samples from our dataset. Where this test set of processed through RootNav2 original and angle based version. It demonstrated that the angle-based method offers lower inferences and processing time for the whole process.

**Table 1.**
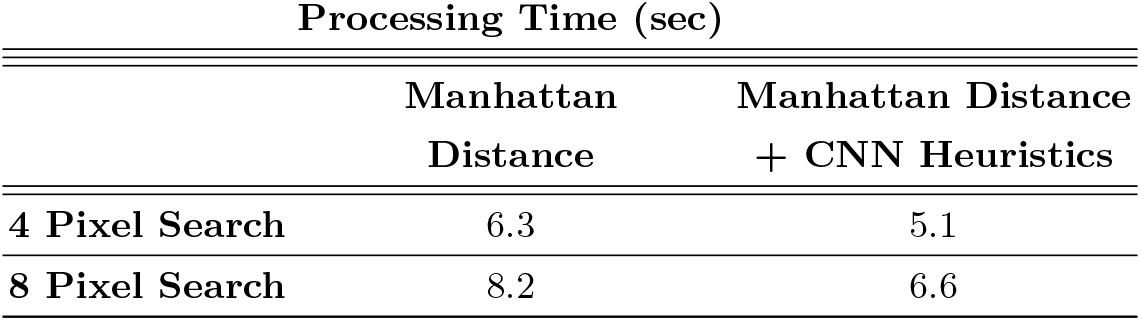
Quantitative Comparison: This graph shows a quantitative comparison of propsoed CNN.

## 5 Conclusions

In this paper, we propose a novel CNN based heuristic function for A* pathfinding algorithm that results in the shortest and smoothest navigation for root system architecture (RSA). It also offers a robots travel from one static root-tip to another (or seed) along a planned path. The experiments have shown that the proposed method offers a far better performance in terms of path search than traditional heuristic functions. It provides low computational cost for roots reconstruction and quality addition to the RootNav 2.0 performance boost. The proposed CNN based heuristic function proved that we could train CNNs as heuristic functions and perform a more intelligent and smart search for pathfinding problems.

## 6 Appendix

### 6.1 Wheat (Triticum aestivum L.) dataset Analysis

The proposed method is transfer learned to Wheat (Triticum aestivum L.) dataset [15]. The results extracted from the proposed method were encouraging and offered very smooth and natural roots outputs. The RootNav Viewer 2.0 used to analyze the results. The Fig. 7 shows output results from Rootnav 2.0 with a heuristic function system. The results are the output of root structure mask and RSML results drawn on original images using RootNav Viewer. It is shown that the output root architecture perfectly fits the original root system and offers a smooth and natural outline of each root structure.

**Fig. 7.**
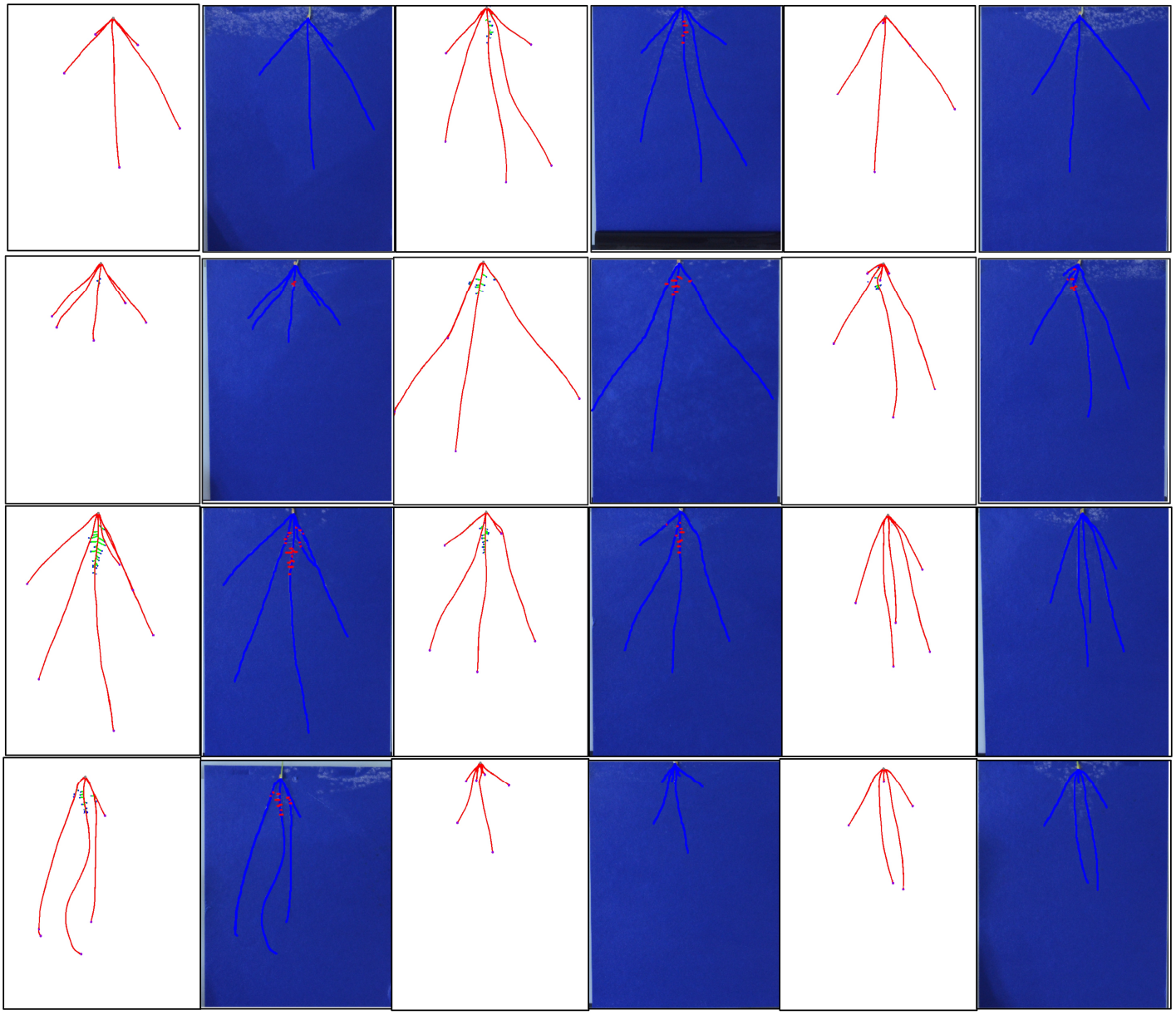
Test Set sample results from RootNav 2.0 + Heuristic function.

### 6.2 Visual Analysis

The Fig. 8 shows a visual analysis of the effectiveness of CNN based heuristic function and its addition to RootNav 2.0. This figure shows how the heuristic function helps the A* algorithm steer along the given root paths and offer more natural-looking roots. The results from Rootnav 2.0 and Rootnav 2.0 with heuristic functions are compared, and it’s clearly shown that later systems in producing much better outcomes.

**Fig. 8.**
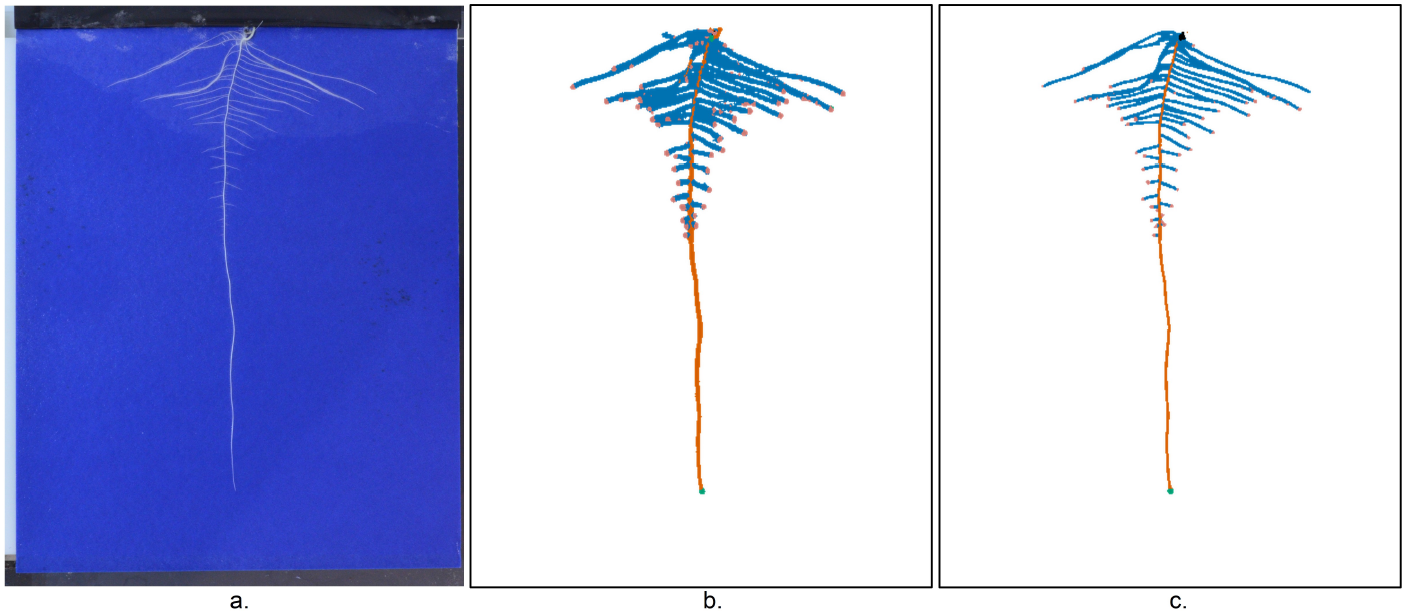
Output Mask Analysis for RootNav 2.0 and RootNav 2 with heuristic function: a. Original input image, b) RootNav 2.0 Colored Mask Output, C) RootNav 2.0+Heuristic-Function Colored Mask Output

### 6.3 Inference Time Analysis

In this analysis, we have used the Test set used in the original RootNav 2.0 [1] research. We have analyzed the original version of RootNav 2.0 with a new angle based AStar algorithm. The results have shown (in Fig. 9) that the new method performs well in the Astar search (primary root search/lateral root search). The overall inference time is also effectively improved, which shows the enhanced performance of the proposed method.

**Fig. 9.**
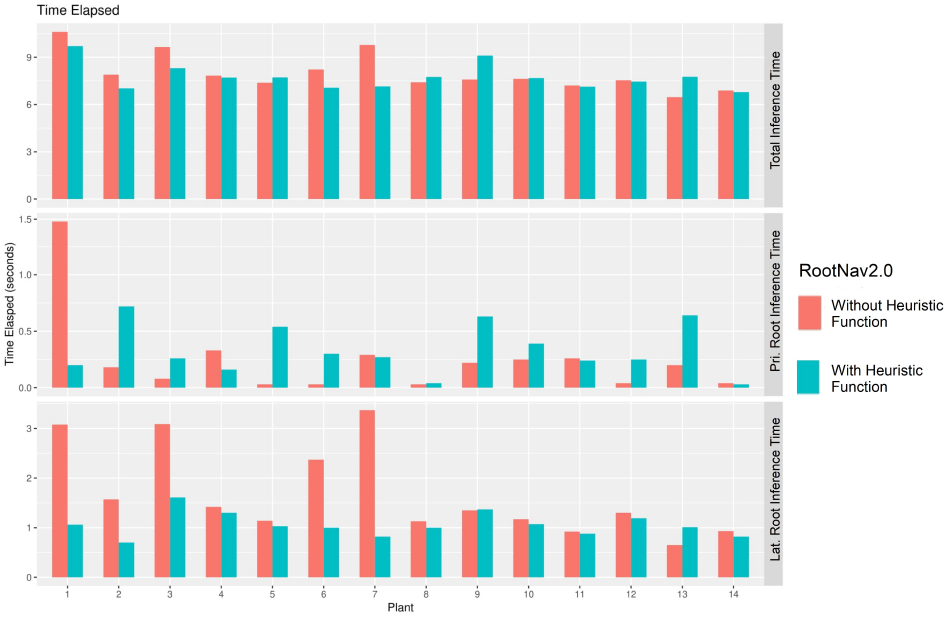
RootNav 2.0 Inference Time Analysis.

### 6.4 Validation of CNN Heuristic Functions

These graphs (Fig. 10) show a graphical overview of heuristic functions based RootNet 2.0 CNN training results for the validation set. The top graph shows the validation Loss and the bottom one shows the proposed system’s validation accuracy. It can be observed that the loss is gradually decreasing, and accuracy is improving alongside, until they become constant. Both curves depict that network trained well, and loss and accuracy converge to the best results naturally.

**Fig. 10.**
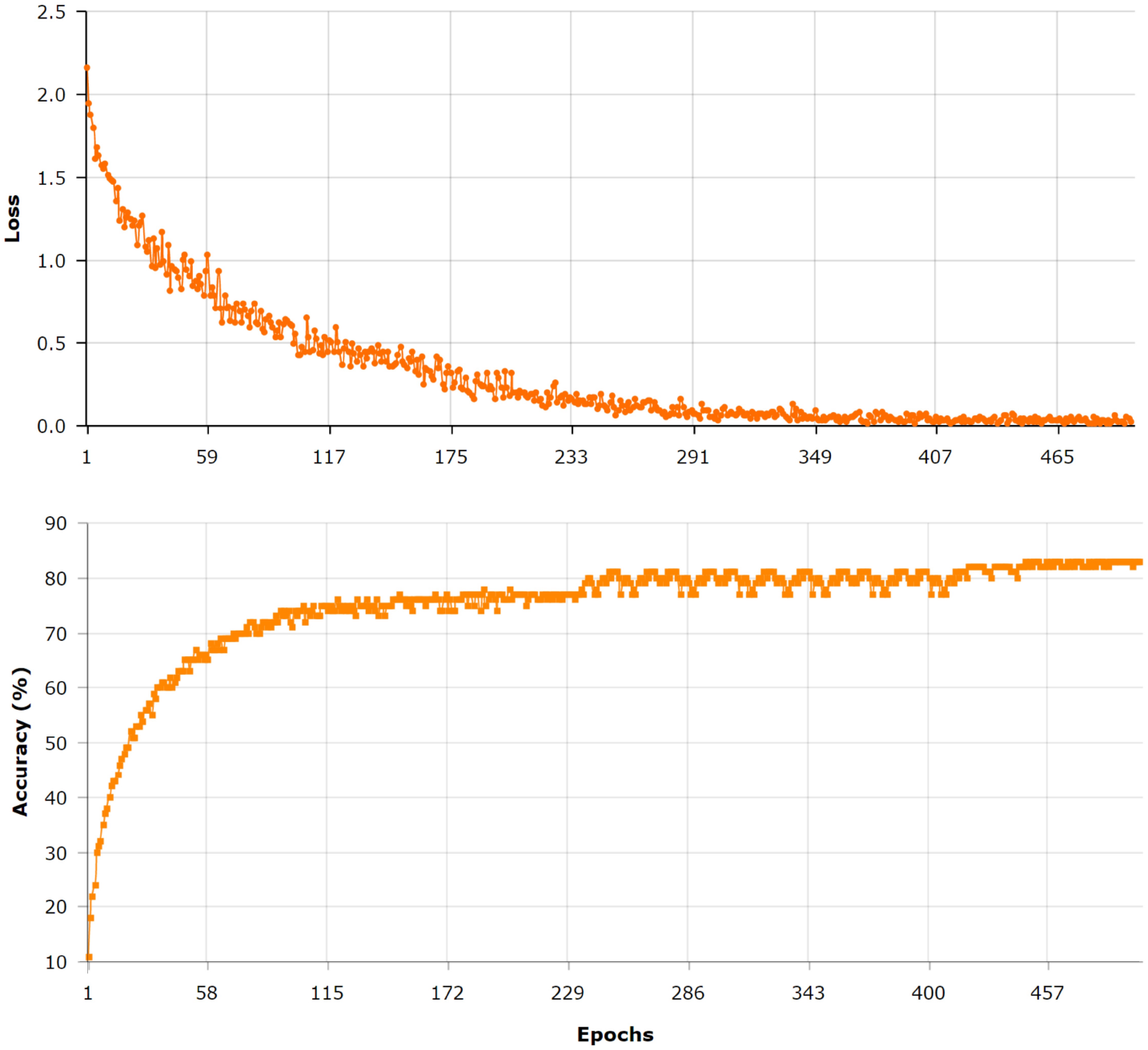
Validation Curves.

